# Design and validation of a multi-point injection technology for MR-guided convection enhanced delivery in the brain

**DOI:** 10.1101/2021.06.24.449788

**Authors:** Kayla Prezelski, Megan Keiser, Joel M. Stein, Timothy H. Lucas, Beverly Davidson, Pedro Gonzalez-Alegre, Flavia Vitale

## Abstract

Convection enhanced delivery (CED) allows direct intracranial administration of neuro-therapeutics. Success of CED relies on specific targeting and broad volume distributions (V_D_). However, to prevent off-target delivery and tissue damage, CED is typically conducted with small cannulas and at low flow rates, which critically limit the maximum achievable V_D_. Furthermore, in applications such as gene therapy requiring injections of large fluid volumes into broad subcortical regions, low flow rates translate into long infusion times and multiple surgical trajectories. The cannula design is a major limiting factor in achieving broad V_D_, while minimizing infusion time and backflow. Here we present and validate a novel multi-point cannula specifically designed to optimize distribution and delivery time in MR-guided intracranial CED of gene-based therapeutics. First, we evaluated the compatibility of our cannula with MRI and common viral vectors for gene therapy. Then, we conducted CED tests in agarose brain phantoms and benchmarked the results against single-needle delivery. 3T MRI in brain phantoms revealed minimal susceptibility-induced artifacts, comparable to the device dimensions. Benchtop CED of adeno-associated virus demonstrated no viral loss or inactivation. CED in agarose brain phantoms at 3, 6, and 9 μL/min showed >3x increase in volume distribution and 60% time reduction compared to single-needle delivery. This study confirms the validity of a multi-point delivery approach for improving infusate distribution at clinically-compatible timescales and supports the feasibility of our novel cannula design for advancing safety and efficacy of MR-guided CED to the central nervous system.

## 1. Introduction

Neurological disorders affect over 100 million people in the United States and pose a significant societal and economic burden, costing more than $800 billion/year in the U.S. alone [1]. The most prevalent and costly are neurodegenerative disorders (NDs) of the central nervous system (CNS) such as Alzheimer’s (AD) and Parkinson’s disease (PD), which affect 6 M people in the U.S., and approximately 42 M people globally [1], [2]. The standard of care for NDs of the CNS are symptomatic pharmacological therapies based on systemic delivery of large molecular weight (MW) drugs administered either orally or intravenously. The blood brain barrier (BBB), however, prevents most of the molecules from entering the interstitium, which significantly hampers the effectiveness of systemic delivery methods.

In recent years, therapeutic development for NDs has shifted from optimization of symptomatic therapies to interventions aimed at altering the natural history of the disease. Many of them, including adeno-associated virus (AAV)-based therapies, require access to brain parenchyma [3], [4]. A successful strategy to bypass the BBB and increase delivery efficiency is based on intraparenchymal (IPa) injections directly into the target site in the brain. This technique is called convection enhanced delivery (CED) and relies on the convective flow generated by a positive pressure gradient imposed by a syringe pump to deliver the infusate through a catheter and into the target brain tissue. Compared to bolus injection and diffusion-driven methods, CED has been shown to achieve significantly higher coverage and volume distributions (V_D_) [5] especially for large MW compounds, since convective flow is independent from MW. CED was first introduced in the 1990s by researchers from the U.S. National Institute of Health (NIH) to enhance the delivery efficiency of drugs that could not cross the BBB or were too large to diffuse over long distances [5]. Since then, CED has been successfully used in IPa delivery of a large number of substances, including chemotherapeutics [6], [7], viral vectors [8]–[12], nanocarriers [13], [14], and neurotrophic factors [15].

In the context of NDs of the CNS, direct IPa administration of AAV via CED is the route of choice in many CNS gene therapy trials [3], [16]–[21] due to: (1) minimal biodistribution to peripheral organs, (2) lower doses, and (3) significantly higher transduction efficiency compared to intravenous [22] or intrathecal [23] delivery. Initial multi-center, double-blind clinical trials involving AAV-mediated gene therapy for PD with neurotrophic factors, such as glial-cell-derived neurotrophic factor (GDNF) [24] and neurturin [16] delivered via bilateral injections in the putamen, failed to achieve primary efficacy endpoints. Retrospective analyses [15], [24] pointed at the limited infusate distribution and the sub-optimal cannula design causing low coverage and off-target delivery as the primary determinants of these poor outcomes. In recent phase 1, open-label trials of AAV2-GDNF for PD [19] even after two serial intraputaminal injections in each brain hemisphere, the putaminal coverage was only 26% and moderate or no clinical improvements in motor scores were observed. Similar results have been reported in recent phase 1 open-label trials of AAV2-L-amino acid decarboxylase (AADC) therapy for PD [25], where volume coverage ranged between 21% and 42% after two or more serial trajectories. Limited efficacy of gene therapy for infantile AADC deficiency [26] and failure of phase III clinical trials for IL13-PE38QQR therapy for glioblastoma [27], have also been attributed to poor target coverage and low V_D_.

The main factors affecting CED performance are the infusion flow rate and the catheter design. As CED is governed by the gradient between skull and injection pressures, the choice of the optimal flow rate is a compromise between maximizing V_D_ and avoiding pressure and stress-induced tissue damage. Typical CED flow rates range from 0.1-0.5 μL/min in rodents [28]–[30] and 3-5 μL/min in pre-clinical and clinical studies [31], [32].

In principle, increasing the cannula diameter could allow higher flow rates and lower total infusion times. However, larger cannula diameter and high flow rates induce the formation of a low-resistance pathway along the cannula tract, which causes the infusate to leak along this pathway and away from the target site. This phenomenon, called backflow or reflux, not only can affect treatment efficacy, but can also lead to unwanted toxic effects due to off-target delivery.

In the last few decades, extensive efforts have been dedicated to improving and optimizing the design of delivery cannulas. Besides minimizing the shaft diameter [25], [33], [34], some of the proposed strategies include polymer coatings [35], microfluidic devices [36], microporous hollow fiber catheters [30], coaxial [37] and recessed [38] cannulas. Multi-point designs, such as the arborizing catheter [39] and the indwelling Cleveland Clinic Multiport catheter (CCMC) [40] have also been proposed. Currently, the most adopted design in pre-clinical and clinical trials involving IPa CED is the step-cannula, where a 0.36 mm fused-silica needle extends 5 mm distally from a central shaft (O.D. 1.5 mm) [41], [42]. Although this step-cannula design has allowed reflux-free delivery at flow rates as high as 10 μL/min in rodent brains [42], in non-human primates and human CED, where flow rates are typically 3 μL/min [12], [31], [41], [43] and do not exceed 5 μL/min [32], the volume distribution is still far from being optimal especially in larger brain regions such as the putamen [18], [19]. Ideally, a cannula design which allows delivery of the required infusate volume in a short amount of time, while minimizing the risk of tissue damage and backflow, would be beneficial for achieving optimal distribution, increasing overall target coverage, and minimizing the procedure duration.

In this work, we present a novel, multi-point injection technology (MINT) for IPa CED in MRI. Instead of a single delivery needle, the MINT device consists of three moveable microcannulas specifically designed to optimize volume distribution and coverage in target regions, while minimizing the number of surgical accesses and total infusion time. We validated the feasibility of MINT specifically for MR-guided CED of AAV through volumetric MRI and AAV compatibility tests. Furthermore, we assessed distribution performance and backflow at varying flow rates through CED of trypan blue dye in agarose brain phantoms and benchmarked the results against single-needle CED in the same model.

## 2. Materials and methods

### 2.1 Device design and fabrication

The MINT device consists of a 30 cm long Nitinol shaft with a 3 mm O.D. and 2 mm I.D., terminating in a conical Polyetheretherketone (PEEK) tip equipped with three openings. The lumen of the shaft houses three moveable microcannulas controlled via a pressure-sensitive plunger and a central actuation system housed in the ergonomic handle made with UV-curable acrylonitrile butadiene styrene resin. The handle is equipped with three flow inlet ports that connect to the infusion pump system (**Figure 1A, B**).

**Figure 1.**
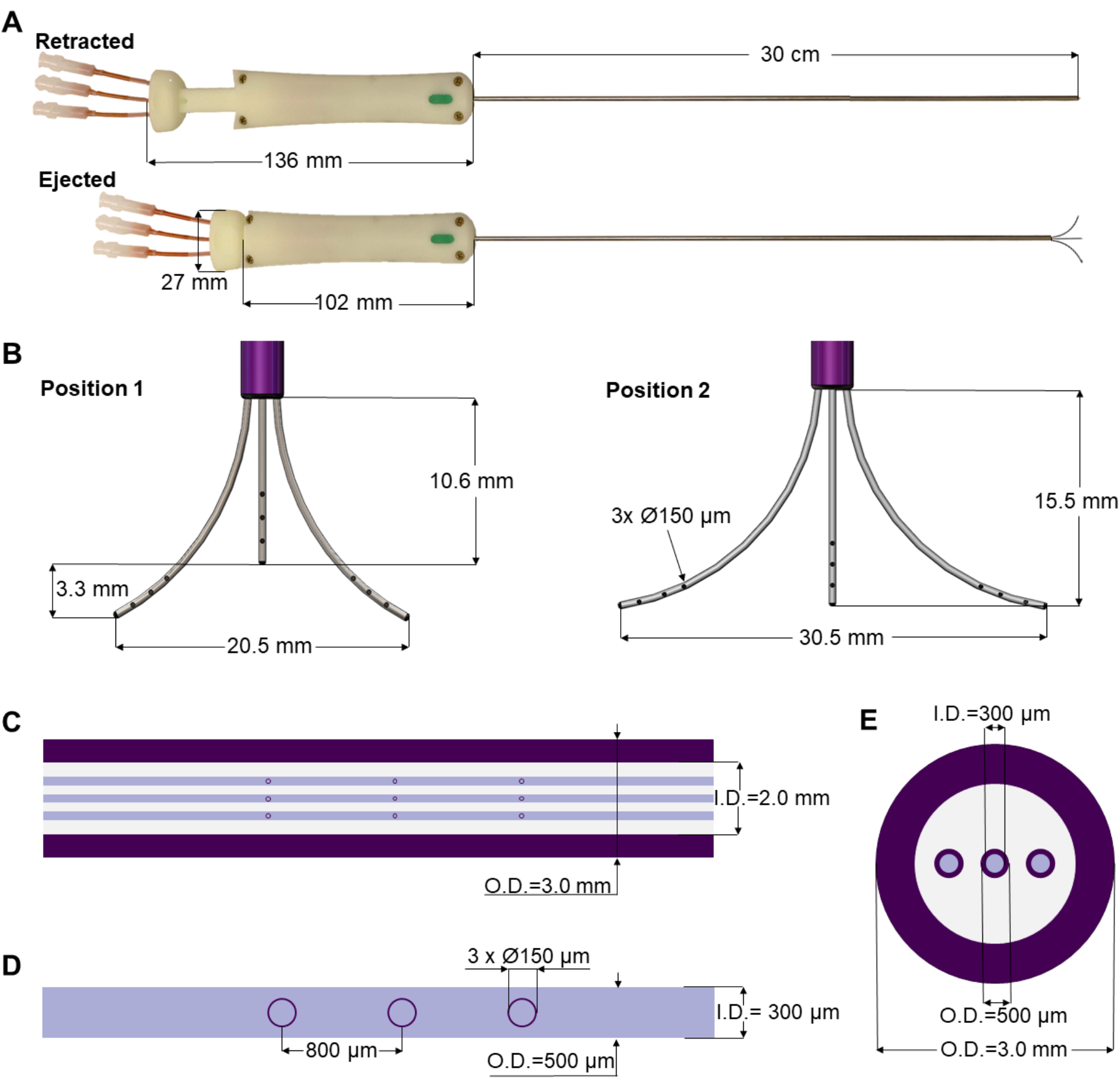
MINT device design and dimensions. (**A-B**) Overview of the dimensions of the MINT catheter showing (**A**) (top) retracted and (bottom) ejected positions, and (**B**) ejected microcannula in (left) intermediately extended Position 1 and (right) fully extended Position 2. (**C-E**) Overview of the dimensions of the three actuated microcannulas including (**C**) details of the shaft dimensions, (**D**) the 3 infusion port dimensions (tapered end not shown), and (**E**) transverse section of the shaft with the embedded microcannulas in the retracted positions.

The microcannulas consist of Nitinol microtubes with 0.5 mm O.D. and 0.3 mm I.D., tapered at the end and machined with three circular fluid outlet points along the distal portion, each 0.15 mm in diameter and spaced 0.8 mm apart (**Figure 1C, D**). Nitinol was chosen because it is an MRI compatible shape memory alloy with exceptional flexibility and resistance against flexural fatigue [44]. The MINT shaft and microcannula diameters are comparable with other single-needle CED cannulas [45]. In addition to the small diameters, the step-design at the transition between the shaft and the microcannulas was chosen as an additional feature to help prevent backflow [28], [39], [42]. The distal ends of the microcannulas are tapered, which has been shown to reduce tissue damage upon insertion compared to blunt tips [46]. The distributed outflow points ensure uniform delivery of the infusate and in previous studies, have been shown to lead to higher V_D_ due to better flow distributions at the outlets [47]. Furthermore, a distributed delivery design leads to lower hydraulic pressure at the fluid outlet, which in turn reduces tissue damage and backflow incidence [30], [47]. The central microcannula is straight, while the two side microcannulas are thermally pre-formed in a curved shape with a maximum radius of curvature of 16.6 mm in the fully extended position. The priming volumes are 46 μL and 49 μL for the central and side microcannulas, respectively.

The microcannula design and curvatures described here were chosen to optimally match the human putamen, a typical target of gene therapies for NDs of the CNS such as PD [18], [19] and Huntington’s disease (HD) [48]. However, these parameters can be easily modified to perform CED in different areas of the human brain as well as for pre-clinical studies in smaller species **(Supplementary Figure 1)**.

### 2.2 MRI compatibility

All the MINT components are made from polymeric materials, except for the Nitinol shaft and microcannulas, and five brass screws and nuts located on the handle. Importantly, there is no closed metallic loop. Nitinol is commonly used in medical devices such as stents and heart valves. Nitinol and brass exhibit less than 10^−3^ susceptibility difference with the brain tissue [49], thus they can be accommodated in the imaging region without causing significant image degradation.

To test the MRI compatibility of the device, we acquired T2-weighted, 3D sampling perfection with application optimized contrasts using different flip angle evolution (SPACE) volumetric images (0.7 × 0.7 × 1 mm) of MINT inserted in a brain imaging phantom prepared according to published protocols [50]. Briefly, the phantom was prepared from 2.9 wt% agarose (IBI Scientific, Dubuque, IA) dispersed in deionized water doped with 21.8 mM NiCl_2_ (Sigma-Aldrich, St. Louis, MO) and NaCl (Fisher Scientific, Hampton, NH). The doped agarose solution was heated and stirred on a hot plate until it became clear, then poured inside a 6-inch, clear-acrylic sphere (American Made Plastic, Inc. Riverside, CA), and degassed in a desiccator. The solution was left to cool and gelate overnight at 4 °C, and then refrigerated until use. The imaging acquisition procedure was performed in a 3T Siemens Trio scanner at the University of Pennsylvania. MRI images were then imported and analyzed offline in OsiriX (Pixmeo SARL, Switzerland).

### 2.3 AAV compatibility

To evaluate whether the materials and fluidic design of MINT are compatible with AAV activity, we conducted AAV compatibility tests in different conditions, ranging from incubation for 30 minutes to CED at varying flow rates. AAV stock solutions at an initial concentration of 1 × 10^13^ vg/mL were prepared according to standard protocols [9], [48]. For incubation tests, the AAV stock solution was manually loaded in one of the MINT microcannulas using a syringe connected to the fluidic line until the microcannula was filled. The AAV solution was incubated for 30 minutes, then collected in microcentrifuge tubes by flushing the line with an air-filled syringe driven by a programmable syringe pump (Harvard Apparatus Inc., Holliston, MA).

Flow tests were conducted by filling the line with fresh AAV stock solutions and running standard CED protocols consisting of an initial stepped flow rate ramping from 0 μL/min to the final flow rate at 0.5 μL/min increments every minute. Finally, 30 minutes of continuous flow at the final flow rate of 3, 5, and 10 μL/min. AAV solution outflowing from the microcannula line was continuously collected in microcentrifuge tubes. At the end of the injection protocol, the remaining AAV solution in the microcannula was collected by manually flushing the line with an air-filled syringe. The concentrations of AAV in the initial stock solution and in each experimental condition were determined by real-time PCR. Briefly, a primer probe set was designed to target a region of the transgene sequence in the AAV used for compatibility testing. A stock standard dilution was created from a linearized plasmid of known size containing the transgene used to generate the virus ranging from 1 × 10^5^ to 1 × 10^11^ copies/mL. The virus used for compatibility testing that was collected under each experimental condition was treated with DNase and diluted 1:1,000, 1:5,000, and 1:25,000 for qPCR analysis. TaqMan master mix (Applied Biosciences Thermofisher Baltics, Vilnius, Lithuania) was used to prepare the qPCR reaction that was run on a CFX384 Real Time Thermal Cycler (BioRad, Hercules, CA). Additional AAV was tested for eGFP transgene expression using an *in vitro* assay. Stock AAVeGFP, AAVeGFP collected after flowing through MINT, or buffer was applied directly to HEK293 cells. HEK293 cells incubated 48 hours before being fluorescently imaged for eGFP positive cells, indicating positive transduction.

### 2.4 CED in agarose brain phantoms: setup

To assess the performance of the MINT device for CED, we developed a shadowgraphy setup and quantitatively measured the volumetric distributions in agarose brain phantoms. The agarose gel was prepared by dissolving 0.6 wt% of agarose (IBI Scientific, Dubuque, IA) into deionized water. The solution was heated and stirred until it became clear and then poured into a custom-made clear-acrylic box (15 cm × 15 cm × 15 cm). The solution was left to cool, gelate overnight at 4 °C, and refrigerated until used. The experimental setup is depicted in Figure 2A and consisted of the clear-acrylic box and a 3D printed top frame designed to securely fit onto the box and rigidly attach to the MRI-compatible stereotactic system SmartFrame (MRI Interventions, Inc., Irvine CA) to guide and adjust the catheter trajectory in the x, y, and z directions. To stabilize the catheter and provide additional support against potential rotation and translation, we used custom, 3D printed reducing tubes and a lateral press-fit post. The three flow inlet ports (**Figure 2B**) on MINT were connected to a programmable syringe pump (Braintree Scientific, Inc., Braintree, MA) via 36” I.V. extension polyethylene tubing (Medline Industries, Inc., Mundelein, IL). Additional components of the setup included a backlight and a side mirror for optimal contrast and accurate reconstruction of the volumetric distribution profiles.

**Figure 2.**
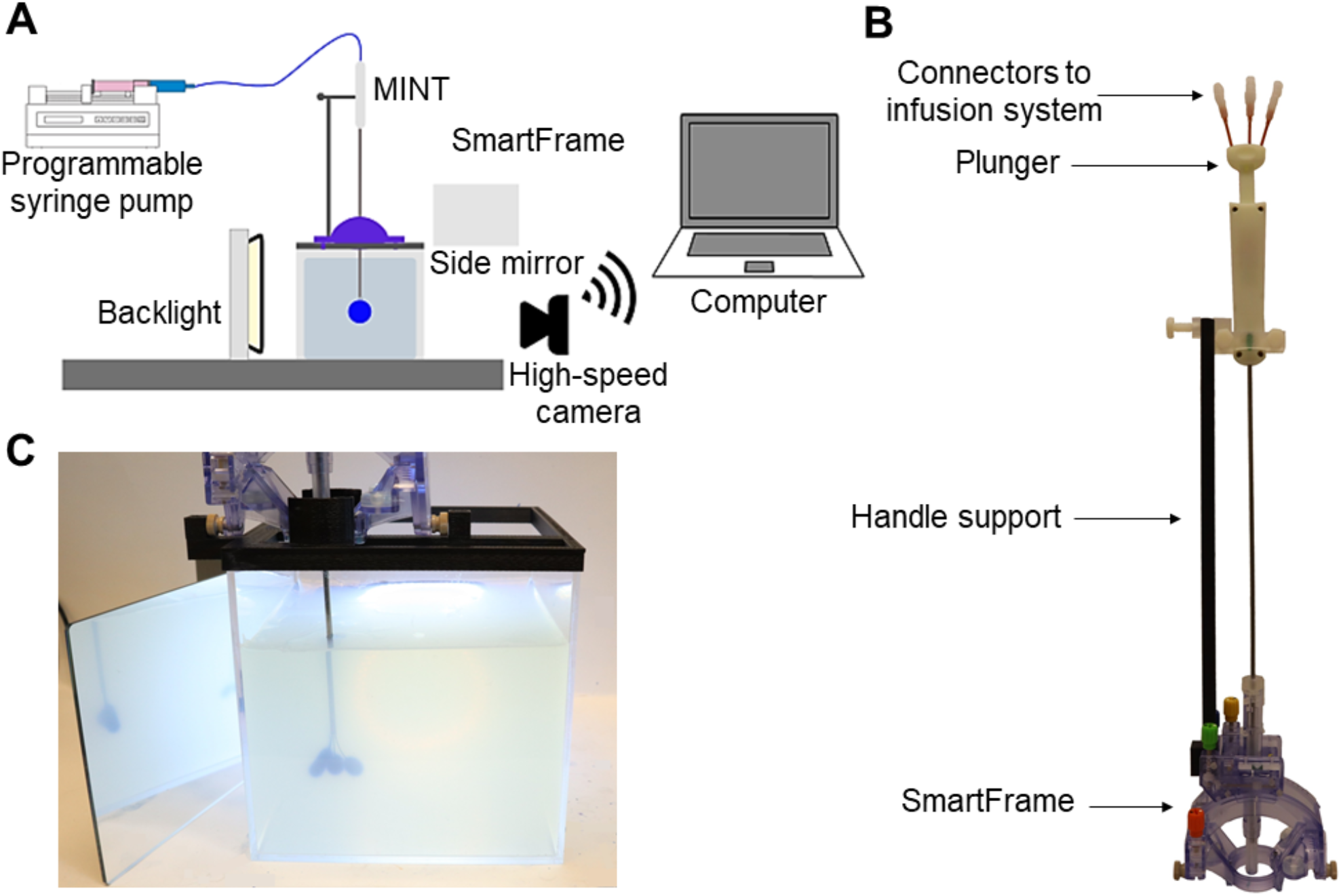
(**A**) Schematics of the experimental setup for phantom CED tests. (**B**) Photograph of the MINT device inserted in the SmartFrame trajectory guidance stereotactic system (MRI Interventions Inc., Irvine CA), with the custom-designed side support. (**C**) Photograph of the MINT device inserted in the agarose brain phantom through the stereotactic frame at the end of the CED of 450 μL of trypan blue dye.

### 2.5 CED in agarose brain phantoms: insertion and infusion protocol

In the CED experiments, the MINT device was connected to the programmable syringe pump. To minimize the risk of the formation of air bubbles and catheter distal tip occlusion during insertion, the pump was turned on at a 0.5 μL/min flow rate before the beginning of the insertion procedure into the agarose gel to maintain a positive pressure.

MINT was then inserted into the stereotactic frame, manually lowered in the agarose phantom, and secured to the lateral support at the desired insertion depth of ~5 cm from the phantom surface. The plunger was then manually actuated at 2.7 ± 1.4 mm/sec to eject the three microcannulas from the shaft (**Figure 2C**). This insertion rate was calculated in Matlab (Mathworks, Inc, Natick, MA) as follows: starting from the last frame of the video, a bulk-crop method was used to select the region of interest containing the central microcannula. The video was then down-sampled from 30 to 6 fps and each cropped frame was converted to a binary image (showing the cannula in white) from which the vertical coordinate of the cannula end was recorded. From these discrete coordinates over the course of the deployment, the microcannula insertion rate was calculated.

Trypan blue dye (0.4%, Sigma-Aldrich, St. Louis, MO) was used as a marker for volumetric flow distribution. The injection procedure followed standard CED protocols [45]: a first ramping step at 0.5 μL/min increments every 1 minute, followed by continuous injection at the desired flow rate. The final total flow rates were set at 3, 6, or 9 μL/min (i.e., 1, 2, or 3 μL/min per microcannula line) and injections were performed until a total volume of 450 μL of trypan blue dye was delivered (**Figure 2C**). This volume is typical for CED infusions of AAV in human gene therapy trials [18], [19]. Images of the volume distribution in the phantom were taken with a Canon EOS M50 4K ultra high-definition digital single-lens reflex camera (Canon, Inc., Ota City, Tokyo, Japan) at 25 μL or 50 μL volume increments. A total of n=3 experiments were performed at each flow rate condition tested.

Following a similar protocol, with steps at 0.5 μL/min increments every 1 minute followed by continuous injection, CED experiments were performed to compare the performance of MINT *Vs.* single-needle cannulas. For these tests, only the MINT central microcannula line was used to mimic single-needle injection. Final flow rates for single-needle experiments were 3 and 5 μL/min.

### 2.6 Volume distribution calculation

The volume distribution is defined as the volume where the injected agent is distributed in the target medium. The injected volume (V_i_) is the volume output by the programmable syringe pump. Therefore, the distribution ratio V_D_/V_i_ is a measure of the infusion efficiency.

The volume distribution over the course of the benchtop infusion experiments was calculated from images of the injected trypan blue dye in the front and side views using the open-source Java-based image processing and analysis software ImageJ. Under the assumption that V_D_ profiles for distributed delivery configurations can be modeled as ellipsoids [30], the semiaxes were manually measured from the trypan blue dye distributions at the end of each microcannula (**Supplementary Figure 2A, B**). The total V_D_ was calculated as the sum of the three ellipsoid volumes:

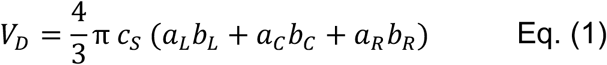

where a, b, and c are the ellipsoid semiaxes and the subscripts L, C, R, and S indicate left, central, right microcannula and side plane, respectively. Statistical analysis was performed using Microsoft Excel with a one-way analysis of variance (ANOVA). A *p-*value < 0.05 was considered statistically significant.

## 3. Results

### 3.1 MRI compatibility

A number of gene therapy protocols for neurological and neurodegenerative disorders involve IPa injections of the viral vectors in the target brain region via MR-guided insertion, CED, and real-time visualization of the infusate distribution [12], [18], [19], [51]. Thus, the MRI compatibility of the CED cannula and lack of susceptibility-induced imaging artifacts are of paramount importance to ensure accuracy of targeting and V_D_ quantification.

Images of the MINT devices in the MRI agarose brain phantom under 3T MRI show the lack of any significant image artifact or distortion (**Figure 3**). Furthermore, we could clearly resolve the position of the three inidvidual microcannulas both in the sagittal and transverse planes, even when they were close to the each other in the partially extended Position 1 (**Figure 1B**). This finding supports the feasibility of detecting volume distribution profiles from each injection cannula during MR-guided CED procedures. The size of the artifact measured on the images was 3 mm in the sagittal plane and 3.3 mm in the transverse plane, which is comparable to the shaft dimensions (O.D. = 3 mm).

**Figure 3.**
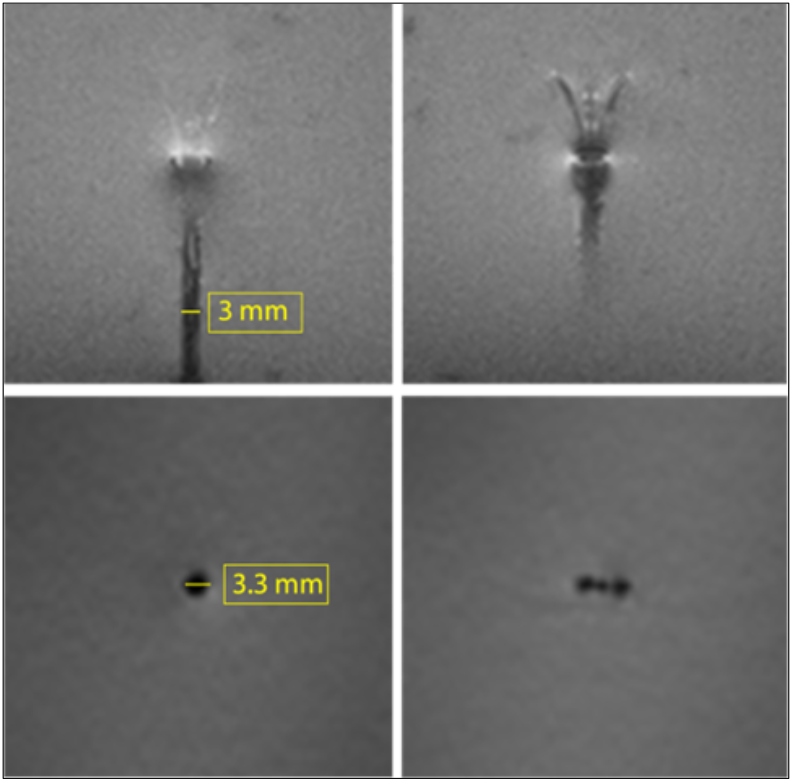
3T MRI of the MINT device in an agarose brain phantom. Top: two consecutive slices in the sagittal plane showing the shaft and the microcannula extended in Position 1 (intermediate extension). Bottom: transverse plane images of the (left) shaft and (right) microcannulas. (Slice thickness:1.3 mm).

### 3.3 AAV compatibility

AAV is one of the most common vectors used for gene therapy in the CNS, due to its low immunogenicity, long-term gene expression profile, high transfection efficiency, and ease of functionalization [3], [17], [20]. When AAV is injected in the target regions via CED, loss or inactivation of AAV can occur via hydrophobic interaction and adhesion to the cannula walls or material toxicity, with potentially detrimental effects on transfection efficiency and therapeutic efficacy [52].

To assess AAV compatibility of the MINT materials and designs, we performed benchtop incubation and CED delivery tests of AAV infusate and quantified the AAV concentration at the end of each tested condition. To simulate a high-velocity, high-shear flow, we also manually flushed AAV infusate from MINT with an air-filled syringe.

Real-time PCR of the AAV collected after 30 min incubation, CED, and air flush conditions did not show loss of AAV compared to the initial titer (**Figure 4**). Furthermore, MINT compatibility analysis conducted on AAV encoding green fluorescent protein (eGFP) showed comparable infectivity in HEK 238 cells to non-exposed vectors (**Supplementary Figure 3**).

**Figure 4.**
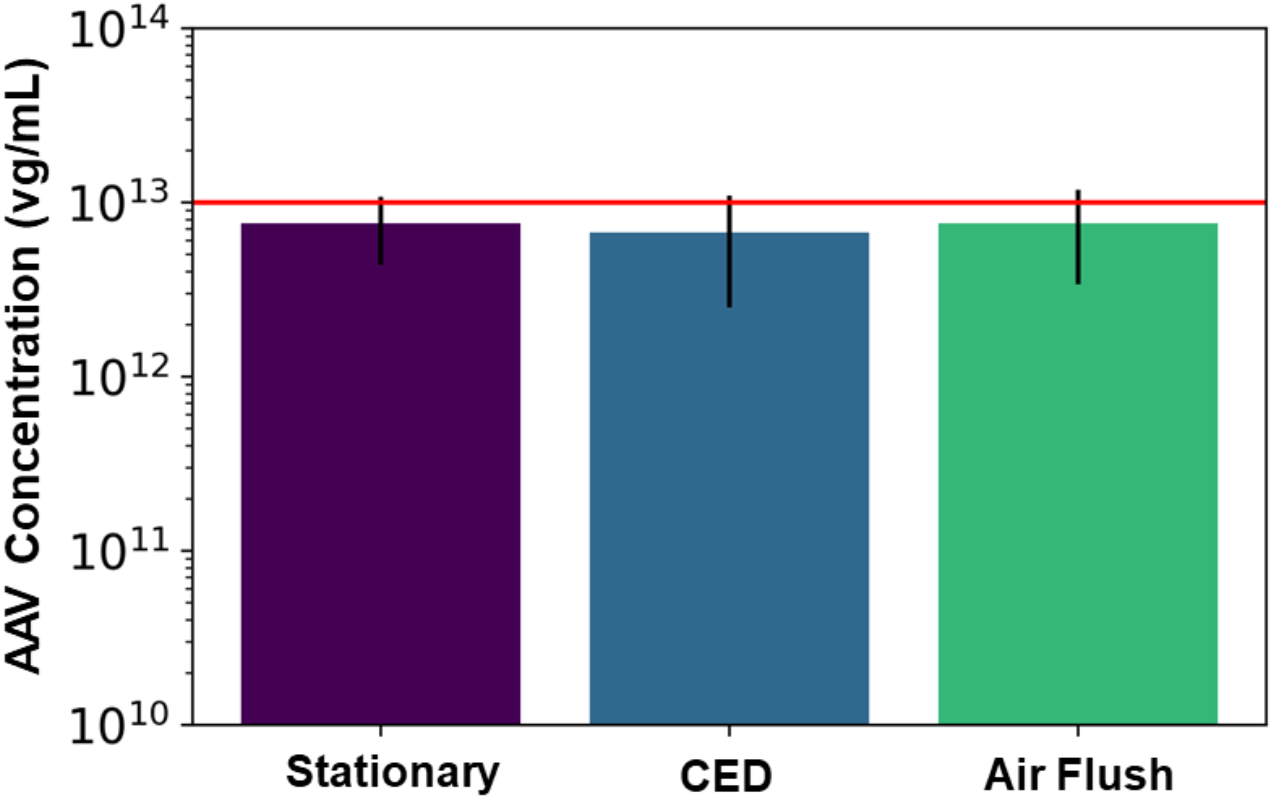
AAV compatibility tests. Bar plots show the concentration of AAV in the fluid collected at the outflow ports in the 30 min incubation, CED, and air flush conditions (n=6 in each condition). Error bars represent ± S.D.

### 3.4 Volume distribution and V_D_/Vi ratio

To assess the performance and evaluate the advantages of our multi-point injection strategy for CED compared to single-needle cannula delivery systems, we simulated the insertion and CED procedure in agarose brain phantoms. Although agarose is inert, non-perfused, homogeneous, and isotropic, previous work demonstrated that 0.6% agarose gels adequately mimic the mechanical properties of brain porous tissue during pressure-driven infusion experiments, resulting in comparable infusate distributions to porcine brain tissue [53].

We conducted our tests at flow rates from each individual microcannula varying from 1, 2, and 3 μL/min (n=3 for each condition). Since we injected the trypan blue dye simultaneously from the three microcannulas, the total flow rates delivered from the MINT device were 3, 6, and 9 μL/min and the total injected volume was 450 μL.

**Figure 5** shows representative trypan blue dye distribution profiles during the course of the experiments at increasing increments of V_i_. Due to the distributed delivery from the multiple outlets along the microcannulas, in all the experiments the infusion cloud morphology showed an ellipsoidal shape uniformly distributed along the distal ends of the microcannulas. Importantly, in all of our experiments, we did not observe reflux of trypan blue dye along the microcannula walls, even at the highest flow rate tested. The reflux-free nature of CED with the MINT device was confirmed by the linear dependence of average volume distribution V_D_ with time (**Figure 6A**) and the constant V_D_/V_i_ profiles after the initial 10-20-minute transients (**Figure 6B**) at all the flow rates.

**Figure 5.**
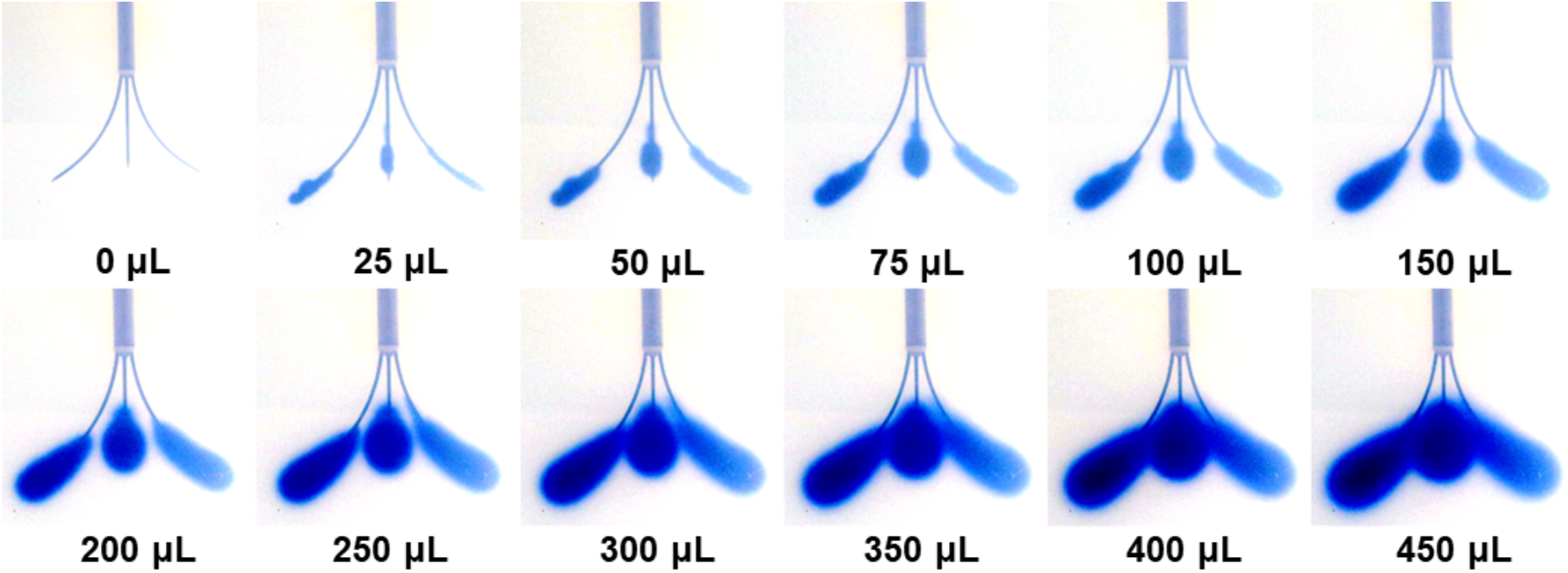
Snapshots of the volume distribution V_D_ during CED injections at 3 μL/min from each microcannula (total flow rate=9 μL/min).

**Figure 6.**
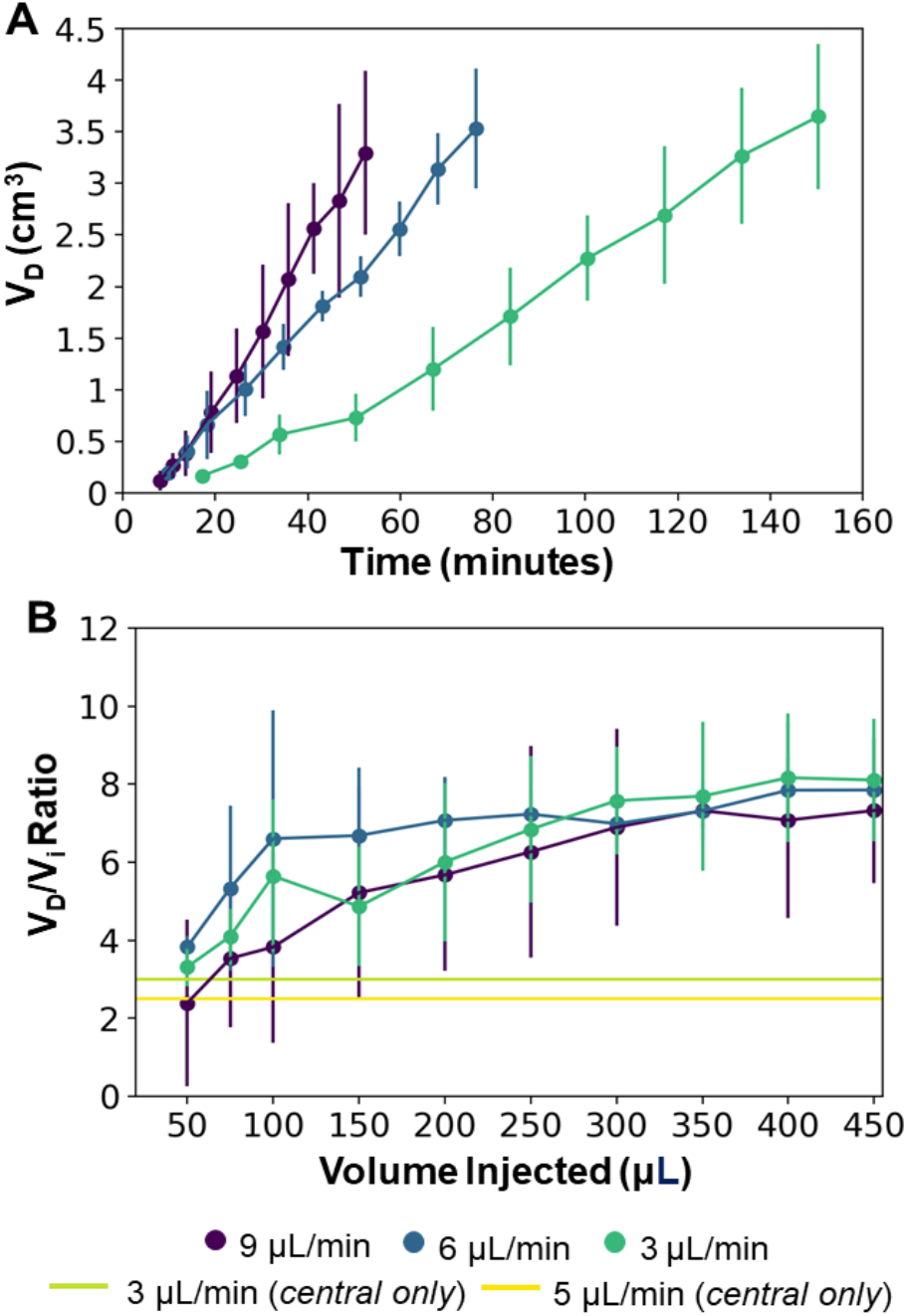
(**A**) Average volume distribution over time and (**B**) average distribution ratio with the MINT devices at varying flow rates. The green and yellow horizontal lines represent the average V_D_/V_i_ in single-needle CED from the central microcannula at 3 and 5 μL/min respectively (data from Supplementary Figure 4). Error bars represent ± S.D. from n=3 trials.

The final values of both V_D_ and V_D_/V_i_ showed a dependence on the total flow rate, although not statistically significant (*p* = 0.82). Specifically, at Vi = 450 μL, V_D_ was 3648.1 ± 704.9 μL, 3533.1 ± 579.3 μL, and 3296.2 ± 792.0 μL, and V_D_/V_i_ was 8.1 ± 1.6, 7.9. ± 1.3, and 7.3. ± 1.9 at flow rates of 3, 6, and 9 μL/min, respectively (**Figures 6 and 7A**). This inverse dependence of the volume distribution from the flow rate is consistent with findings from previous works [36], [37], [54], [55] and can be attributed to the reduction in the permeability of the gel porous matrix caused by the perfusion-induced deformations at higher flow rates (i.e., effective pore size reduction). The total delivery time was 150.4, 76.4, and 52.4 minutes at 3, 6, and 9 μL/min respectively, which is 30% of the time required to deliver the same infusate volume from a single cannula at a given flow rate. At 3 μL/min, the distribution ratio V_D_/V_i_ from the multi-point CED injections with the MINT device was 2.7-fold and 3.2-fold higher than V_D_/V_i_ measured during single-needle CED experiments at flow rates of 3 and 5 μL/min, respectively. Specifically, the average V_D_/V_i_ from single-needle injections were 3.0 ± 0.5 at 3 μL/min and 2.5 ± 0.7 at 5 μL/min (**Supplementary Figure 4**).

**Figure 7.**
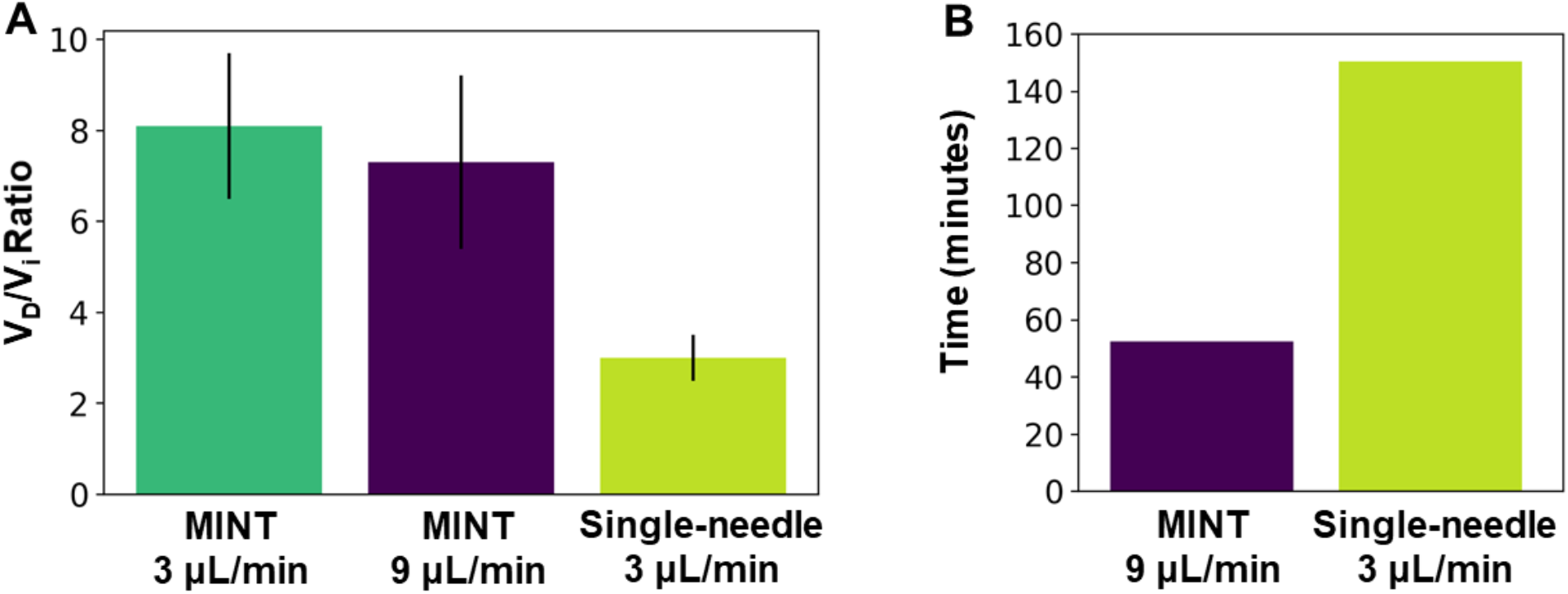
(**A**) Average distribution ratio for multipoint *Vs* single-needle CED. (**B**) Infusion duration for the multipoint *Vs* single-needle CED. Error bars represent ± S.D. from n=3 trials for multipoint at each flow rate condition and n=4 trials for single-needle CED.

## 4. Discussion

In this work, we described and validated a novel multi-point injection design for IPa CED. We engineered our device to achieve maximal volume distribution, while minimizing infusion time, risk of tissue damage, and backflow from large cannula size and elevated flow rates. The key design features of the MINT device to ensure efficient CED and enhanced V_D_ are: (1) three actuated microcannulas for simultaneous multi-point CED and large volume coverage; (2) distributed delivery points along the microcannulas to reduce outlet pressure and achieve uniform infusate distribution; (3) step-design, tapered microcannula ends, and small diameters to reduce risk of tissue damage and backflow occurrence.

Furthermore, as delivery is a critical factor in accelerating the translation of gene therapy into clinical care for the treatment of NDs of the CNS, we specifically chose materials and design features that allow MINT to readily integrate into standard platforms for MR-guided CED of viral vectors via the IPa route.

Typically, IPa CED procedures are conducted in an MRI scanner for real-time monitoring of catheter placement and agent coverage [18], [19], [56]. Our MR-compatibility tests show minimal susceptibility-induced artifacts which are comparable with the device dimensions in standard volumetric 3T MRI sequences. Furthermore, the size and geometric features of the MINT device make it possible to readily integrate MINT within standard stereotactic apparatuses for MR-guided CED procedures, such as the SmartFrame (MRI Interventions, Inc., Irvine CA). These results support the feasibility of integrating MINT into current MR-guided insertion and injection neurosurgical protocols, and to accurately detect cannula targeting and volume distribution profiles during the procedures, in real-time.

AAV compatibility of a CED device is particularly relevant in application of CNS gene therapy, where AAV is the most common vector of choice. Thus, loss of AAV infectivity coupled with low volume coverage could be detrimental for the therapeutic efficacy and translational potential of gene therapy platforms. Benchtop injection tests on AAV articles at clinically relevant flow rate conditions show that the chemical, physical, and design properties of the cannula are compatible with AAV and do not cause virus loss or inactivation of the infectivity.

To assess whether our novel multi-point injection cannula design resulted in improved volume distribution performance compared to the single-needle design, we conducted CED tests of a tracer dye in agarose brain phantoms. This study revealed that simultaneous infusions through three microcannulas with a multiple-opening design result in ~3x higher volume distribution compared to single-needle CED (**Figure 7A**). The advantage of our multi-point cannula configuration is also evident when compared with other single-needle CED cannulas: in agarose brain phantoms the distribution ratio V_D_/V_i_ with MINT is 23% higher than the Valve Tip (VT) catheter (Engineering Resources Group), and 50% higher than the SmartFlow step-design single-needle cannula (MRI Interventions, Inc., Irvine CA) (**Table 1**) [45]. The MINT distribution ratio falls only 42% lower than a research-grade multi-port arborizing catheter with seven delivery cannulas, compared to MINT with three microcannulas [39]. Other multi-cannula geometries have been proposed in literature, such as the Cleveland Clinic Multiport catheter [40], but they have been designed as indwelling devices for continuous, multi-day injections (96 hs) and thus, distribution data cannot be directly compared.

**Table 1.** Comparison of V_D_ for different single-needle and multi-point CED cannulas

| Device | $V_D/V_i$ | Flow Rate | End-point Type |
| --- | --- | --- | --- |
| VT Catheter [45] | 6.6 | 3 $\mu\text{L}/\text{min}$ | Single |
| SmartFlow Catheter [45] | $5.4 \pm 2.3$ | 3 $\mu\text{L}/\text{min}$ | Single |
| Arborizing Catheter [39] | 14.9 | 7 $\mu\text{L}/\text{min}$ | Multi (7) |
| MINT | $8.1 \pm 1.6$ | 3 $\mu\text{L}/\text{min}$ | Multi (3) |

A larger distribution ratio V_D_/V_i_ is indicative of improved coverage of the target region, a main factor determining the ultimate outcomes of therapeutic paradigms relying on direct IPa delivery. For example, AAV transfection efficiency has been shown to directly correlate with 3D volume distribution measured during MR-guided CED [43], [57]. In many gene therapy protocols for NDs of the CNS the target structure is the putamen, an irregularly-shaped, large volume, subcortical nucleus (3.6 cm^3^ in humans [58]). Although current intraputaminal CED delivery protocols require two or more surgical trajectories and 450 to 900 μL injections of AAV articles in each hemisphere [18], [19], typical putaminal volume coverages range between 21% and 42% at best [18], [19], which is well below the minimum target of 60% [59]. In our brain phantom experiments the volume distribution from 450 μL injections ranged between 3.6 and 3.3 cm^3^, which is higher than the target minimum coverage in the human putamen. Although the transport properties of the putaminal tissue are different than those of the agarose gel – due to perfusion, anisotropy, and non-homogeneities – and, thus, these results cannot be directly extrapolated to predict putaminal coverages, they support the feasibility of the proposed multi-point injection device to improve distribution in CED.

An additional and relevant advantage of our approach is the significant reduction in total infusion time (**Figure 7B**). The long duration of the CED procedures makes IPa particularly risky and challenging not only in large structures such as the putamen, but also in less accessible regions such as the cerebellum, a target for gene therapy of different forms of spinocerebellar ataxias [8], [9], [60]. By multiplexing the CED protocol simultaneously across multiple infusion sites, the MINT device can deliver 3x the infusate volume of a single-needle cannula in a given amount of time while operating at clinically relevant flow rates (3-9 μL/min). Thus, MINT enables a significant reduction in the duration of the injection procedure without impacting the delivery performance.

Finally, although maximizing volume distribution was the primary objective of this work, the step change at the interface between the main shaft and the microcannulas, together with the tapered end profiles and small diameter, helped to minimize the occurrence of backflow during the infusion procedures conducted in our study.

The rate of cannula insertion has been shown to be another factor affecting backflow incidence [54]. Although in many previous works the experimental protocols involved gelation of the agar phantoms around the cannula [37], [39], in our study we included the manual catheter insertion and microcannula deployment steps to more realistically mimic actual CED procedures. Future work will be devoted to investigating the effect of varying insertion and deployment rates on backflow, with the goal of defining the optimal rate parameters for the multi-point CED procedure.

Additional future experiments will include comparative analysis of volume distributions in non-homogeneous substrates, such as animal and eventually, human brain tissues. Particularly relevant to validate our technology against the anatomical and transport challenges of the brain parenchyma will be *in vivo,* pre-clinical MR-guided targeting, and co-injections of contrast agents, such as gadolinium, and viral vectors tagged with fluorescent reporters, to accurately track distribution, coverage, and transfection efficiency via volumetric MRI analysis and *post-mortem* histology, respectively [48], [61].

## 5. Conclusion

Direct drug and gene delivery to the brain has the potential to become a truly curative therapeutic option for those affected by neurological disorders by circumventing the challenges of the blood-brain barrier penetration. In order to fully realize this potential, the issue of targeted and broad infusate distribution via CED must be addressed via novel engineering solutions to the delivery cannula systems.

In this work, we have proposed and validated MINT, a novel multi-point injection cannula for achieving broader volume distribution than current single-needle designs. We have validated our system in benchtops studies of trypan blue CED in agarose brain phantoms, demonstrating significant increase in volume distribution compared to single-needle delivery, while drastically reducing the total duration of the procedure. Furthermore, we have demonstrated that our device is compatible with both the most advanced protocols for MR-guided insertions and injections and viral vectors used in a number of CNS gene therapy platforms.

Overall, this study supports the feasibility and the translational potential of a multi-point injection approach as a potentially transformative and enabling solution for highly efficient CED delivery of gene-based therapeutics in the brain.

## Supporting information

Supplementary Infromation

## CRediT statement

K.P. Methodology, formal analysis, software, visualization, writing – original draft. M. K. Data curation, methodology, writing – review & editing. J. M. S., T. H. L., B. D. Resources, supervision, writing – review & editing. P.G.A Conceptualization, resources, supervision, funding acquisition, writing - review & editing. F.V. Conceptualization, resources, supervision, funding acquisition, project administration, writing – original draft.

## Acknowledgments

This work was supported by the NIH R01NS117756 (F.V. and T.H.L) Penn Center for Health, Devices and Technology (F.V. and P.G-A.), the Mirowski Family Foundation, and Neil and Barbara Smit (F.V.).

