## Supplementary Infromation for "Design and validation of a multi-point injection technology for MR-guided convection enhanced delivery in the brain"

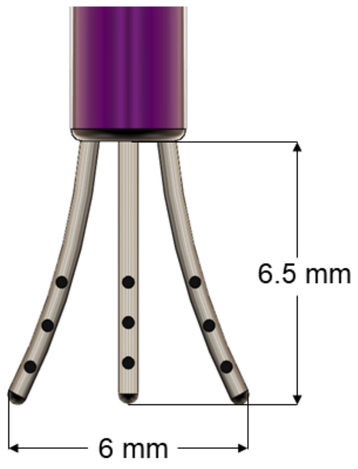


**Supplementary Figure 1.** Overview of the dimensions of the MINT catheter showing the ejected microcannula position of the device designed for infusions in non-human primates.

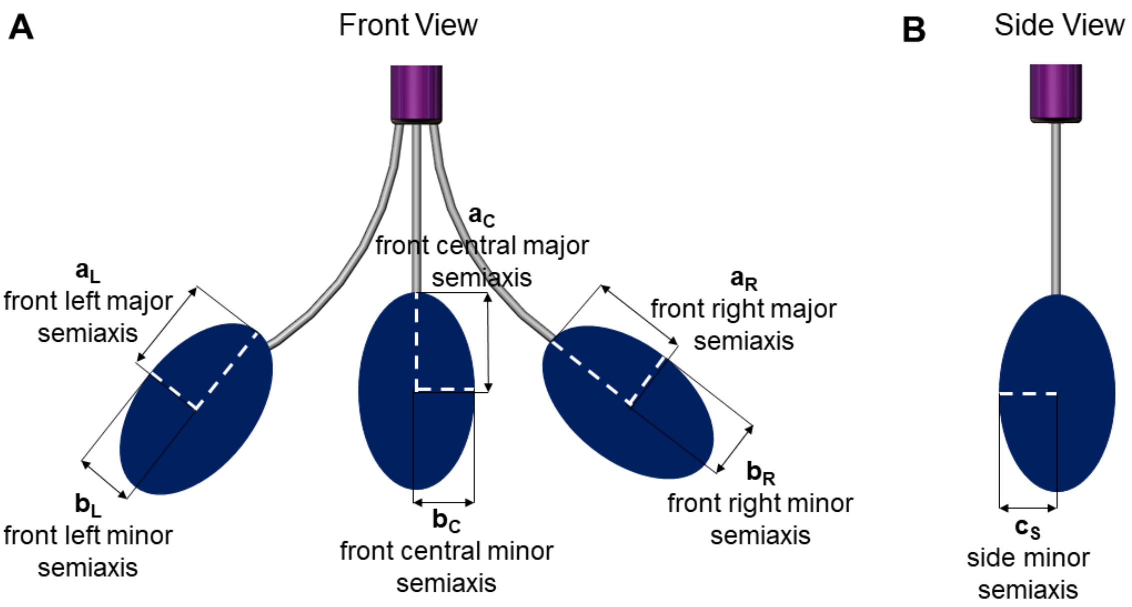


**Supplementary Figure 2.** Volume distribution was calculated as the sum of the volume of the three ellipsoids according to the formula $V=4/3\pi abc$. (**A**) The major and minor semiaxes of the front view and (**B**) the minor semiaxis of the side (mirror) view were used in the calculations as shown.

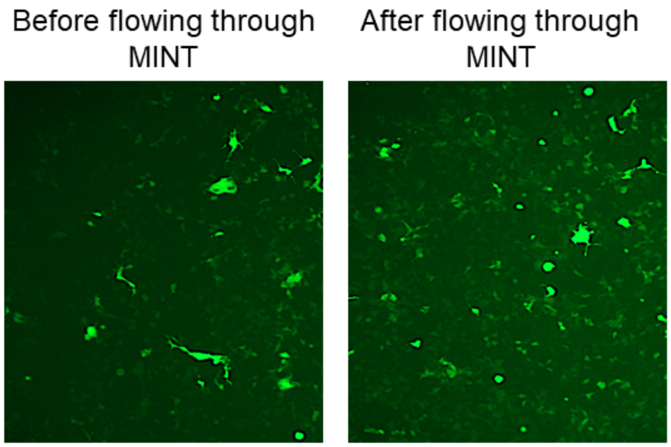


**Supplementary Figure 3.** Virus activity assay in HEK293 at 48 hours post transfection. eGFP fluorescence marks transfected cells from AAV (left) not exposed to MINT, (right) after flow in MINT.

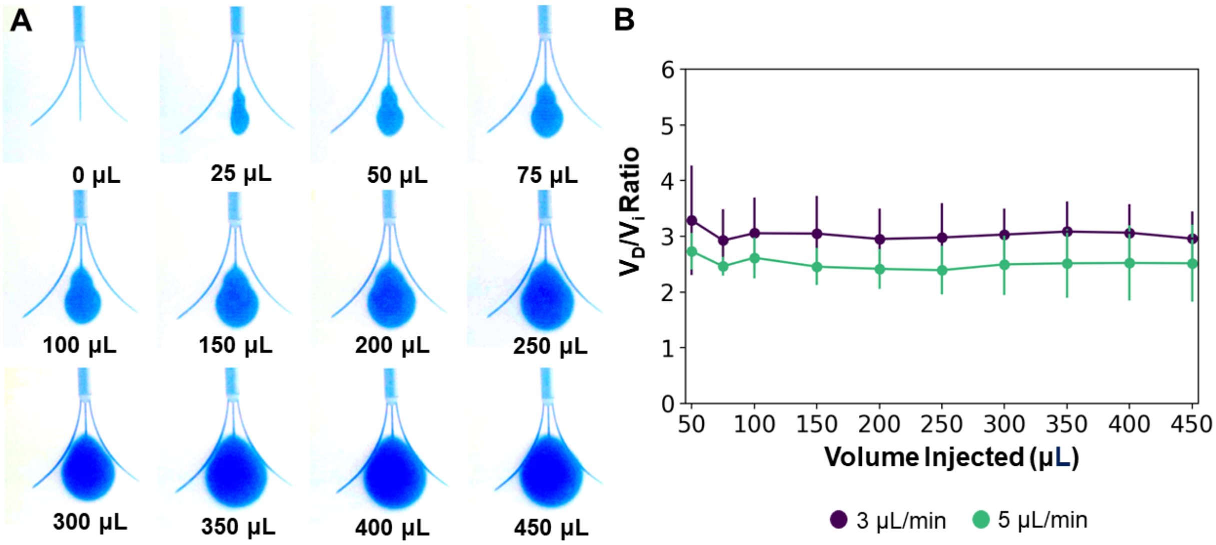


**Supplementary Figure 4.** (**A**) Snapshots of the volume distribution $V_D$ during single point CED injections from only the MINT central microcannula. (**B**) Average $V_D/V_i$ at from the MINT central microcannula at 3 and 5 µL/min total flow rate. Error bars represent S.D. for n=4 trials.
